# *De-novo* draft genome sequence of *Crocus Sativus* L, Saffron, a golden condiment

**DOI:** 10.1101/2021.06.23.449592

**Authors:** Sheetal Ambardar, Jyoti Vakhlu, Ramanathan Sowdhamini

## Abstract

*Crocus sativus* L, saffron is the highest priced but low yielding plant of medicinal and culinary importance. Despite its economic status, the omic information on this plant is very scarce, with only a couple of transcriptomics and epigenetic studies. In the present study, the draft genome sequence of *C. sativus* has been assembled using Illumina sequencing. In fact, this is the first genome sequence from any member of family *Iridaceae*. Genome size of *C. sativus* was estimated to be 3.5 Gb and the draft genome is 3.01 Gb long with 84.24% genome coverage. In total, 8,62,275 repeats and 9,64,231 SSR markers have been identified. A total of 53,546 functional genes were annotated, out of which, 43,649 proteins were associated with GO annotation. 5726 proteins were identified as transcription factors, with MYB & MYB related family proteins being more abundant. Orthology analysis of *C. sativus* with 3 different monocot species of the same plant order and rice (model monocot plant) revealed 7328 proteins clusters to be conserved in all the five plant species, whereas 2510 proteins cluster were unique to *C. sativus* only. 10,912 unigenes of *C. sativus* were mapped to 387 KEGG pathways of monocot. The genes involved in the pathway of apocarotenoids biosynthesis (crocin, crocetin, picrocrocin, and safranal) were present in the draft genome.

## Introduction

Plant genomics, with the increasing number of whole genome sequences available, has unlocked the genetic treasures, that would be impossible in absense of the genome sequence. Though second and third generation sequencing technologies, coupled with ever advancing bioinformatic tools/pipelines, have made the sequencing of complex and huge genomes economical and easy, but still till date there are only approximately 1565 plant genome sequenes available in databanks (NCBI:https://www.ncbi.nlm.nih.gov/assembly). Some of the recently sequenced and assembled plant genome are rice (Choi et al., 2020), Maize (Liu et al., 2020), Asparagus (Harkess et al., 2017), Wheat (Alonge et al., 2020) and tea (Xia et al., 2020) etc., however the genome of the plants belonging to Crocus genera or *Iridaceae* family, have not reported so far.

Saffron (*C. sativus*) referred as ‘Golden Condiment’ is world’s most expensive spice costing about 70,000 INR/pound, with medicinal properties and cosmetic uses (Mzabri et al., 2019; Magotra et al., 2021). More than 150 volatile and aroma-yielding compounds contribute to the flavor, color, and aroma of the Saffron spice, wherein the main chemical constituents in the stigma of saffron are crocin, crocetin, picrocrocin, and safranal (Samarghandian et al., 2014; Maggi et al., 2020). *C. sativus* is an autumn-flowering perennial sterile triploid plant (2n = 24) with, ~3.5 Gb haploid genome (Brandizzi and Caiola 1996, 1998). Being sterile, it fails to produce viable seeds and reproduces vegetatively by underground corms and is reported to lack genetic variation. Various molecular markers (RAPD, ISSR, AFLP, SSR microsatellites) and epigenetic approaches have suggested the existence of limited genetic variability (Rubio-Moraga et al., 2009; Siracusa et al., 2013; Busconi et al., 2018; Mir et al., 2021). To discover authentic genetic markers, mining genes for secondary metabolites and improvement of breeding, sequencing of its genome was the only alternative. In addition, it’s ancestry is also controversial (Alsayied et al., 2015; Nemati et al., 2019), that could be also settled, if its complete genome sequence is available.

Hybrid sequencing approaches, comprising of second and third generation sequencing technologies, have facilitated sequencing of complex genomes economically. Illumina sequencing technology is preffered for first sequencing attempt, as it generates good sequencing data for better genome coverage and has low error rate as compared to third generation sequencing technologies (Edwards and Batley 2010). In the present study, *C. sativus de novo* draft genome sequence has been assembled using short read Illumina sequencing technology. The draft genome sequence unravels the putative markers, metabolic pathways, transcription factors and orthologous genes.

## Materials and Methods

### Sample collection, genome size estimation, whole genome library preparation and Sequencing

*C. sativus* corms were collected from Kishtwar, J&K (33.3116° N, 75.7662° E) in 2019. Corms were grown in the pots for period of three months and leaves were harvested for genome size estimation. Genome size of the plant was estimated by flow cytometric (Hare and Johnston 2011) and k-mer based method using JellyFish (Marçais and kingsford 2011). Genomic DNA was extracted from corm tissue using CTAB method (Rogers and Bendich 1994) and quality and quantity was accessed using Qubit (Invitrogen) and agarose gel electrophoresis. 3 ug DNA was used to construct WGS DNA libraries with 550bp and 800bp insert sizes using NEB next Ultra DNA Library Preparation Kit according to the Illumina’s protocol. Quality of the libraries were evaluated using Tapestation (Agilent 4200) and Qubit HS DNA Assay Kit (Invitrogen). Sequencing was done on the HiSeqX platform (150-bp paired-end (PE) reads) to generate 321 Gb data (~92X coverage).

### *De-novo* Genome Assembly and statistics

Quality of raw reads was evaluated using FastQC tool (Andrews S. 2010) and low quality bases (<q30) and sequencing adapters were removed using trimmomatic software (Bolger et al., 2014). De-novo genome assembly was performed using two assemblers namely Soapdenovo2 (Luo et al., 2012) and MaSuRCA (Zimin et al., 2013). Soapdenovo2 assembly was executed using different kmers (73 kmer predicted by KmerGenei along with 69, 71 kmers) (Chikhi and Madvedev 2013). The statistics of soapdenovo2 assemblies were compared to select the better assembly that was designated as Cs_Assembly_1. MaSuRCA assembly was done using the raw reads and was designated as Cs_Assembly_2. The quality of assemblies was accessed using BUSCO against Viridiplantae lineage from OrthoDB database (Simao et al., 2015). Subsequently, raw illumina reads were mapped back to Cs_Assembly_2 using Bowtie2 (Langmead 2010) and previously published transcriptome data (Baba et al., 2015; Jain et al., 2016) and mapped to Cs_Assembly_2 using BWA (Li and Durbin 2010). Repetitive regions in Cs_Assembly_2 was identified using Repeatmasker and GenomeScope v2 (Ranallo-Benavidez et al., 2020, Smit et al., 2015) and SSR markers were identified using MISA (Beier et al., 2017).

### Genome prediction and annotations, orthology and metabolic pathway analysis

Cs_Assembly_2 was further analysed for gene prediction using the MAKER (Campbell et al., 2014) wherein *C. sativus* transcriptome data was used as EST evidence (Jain and coworker 2016), Viridiplantae database (UNIPROT) as protein evidence, maize as Augustus gene prediction model and *Oryza sativa* as snap hmm. Predicted proteins were further annotated using BLASTp against NR (NCBI) and viridiplanteae (UNIPROT) database with modified parameters (E-value-1e^-3^, sequence identity >40% and query coverage >70%). Annotated proteins were analysed for GO annotations against biological processes, cellular component and metabolic processes using WEGO (Jia et al., 2018). Transcription factors (Tfs) proteins were identified against PlantTFDB (Jinpu et al.,2017) using BLASTp with the modified parameters (E-value-1e^-3^, sequence identity >30%, query coverage >70%). Orthologous genes were compared with *Asparagus officinalis, Dendrobium catenatum, Phalaenopsis equestris, Apostatia shenzhenica* of the same plant order along with *Oryza sativus* (Rice) using Orthovenn2 (Xu et al., 2019) The proteins sequences of all the plants were downloaded from Phytozome database (David et al., 2012). Various metabolic pathways in *C. sativus* genome were analysed using KAAS webserver (Kyoto Encyclopedia of Genes and Genomes Automatic Annotation Server) (Moriya et al., 2007).

### Data availability

Whole genome sequencing raw reads has been submitted to NCBI SRA under bioproject PRJNA734464. The draft genome of *Crocus sativus* has been submitted in SRA under bioproject PRJNA739096. All the processed data including draft genome, annotated proteins, SSR, Transcription factors, and supplementary tables and figures can be accessed at http://caps.ncbs.res.in/download/csat.

## Results and Discussion

*Crocus sativus* genome, being reported here, is the first draft genome sequence of the plant belonging to the *Iridaceae* family. Genome size of *C. sativus* was estimated to be 3.5 Gb (3,578,575,507 bases), using flow cytometry and kmer method. Genome size estimated was comparable to earlier reports, wherein it was estimated to be 3.44 Gb using flow cytometry being grown in Italy, Spain and Israel (Brandizzi and Cajola 1997; 1998). On the basis of size of the genome, 321 Gb WGS data of *C. sativus* was generated, with an overall coverage of ~92X using Illumina sequencing (Table 1). *De novo* genome assembly and annotation of *C. sativus* was performed using the bioinformatics pipeline represented in Fig 1 for easy comprehension.

**Table 1:** Total raw data of 321.36 Gigabases obtained from two insert size (500 bp and 800 bp) libraries using Illumina sequencing with an overall coverage of ~92X. The genome size was estimated as 3.5 Gigabases (3,578,575,507 bases).

| **Libraries**<br>**insert Size** | **Sample Name** | **%**<br>**GC** | **Sequences**<br>**(Million**<br>**Read)** | **Data**<br>**(in Giga-**<br>**bases)** |
| --- | --- | --- | --- | --- |
| 800 | Lib1_800_R1 | 46% | 93.2 | 13.98 |
|  | Lib1_800_R2 | 46% | 93.2 | 13.98 |
|  | Lib2_800_R1 | 46% | 357.8 | 53.68 |
|  | Lib2_800_R2 | 46% | 357.8 | 53.68 |
| 500 | Lib3_500_R1 | 46% | 300.1 | 45.02 |
|  | Lib3_500_R2 | 46% | 300.1 | 45.02 |
|  | Lib4_500_R1 | 46% | 320.0 | 48.01 |
|  | Lib4_500_R2 | 46% | 320.0 | 48.01 |
| **Total** |  |  | **2142.2** | **321.36** |

**Fig 1:**
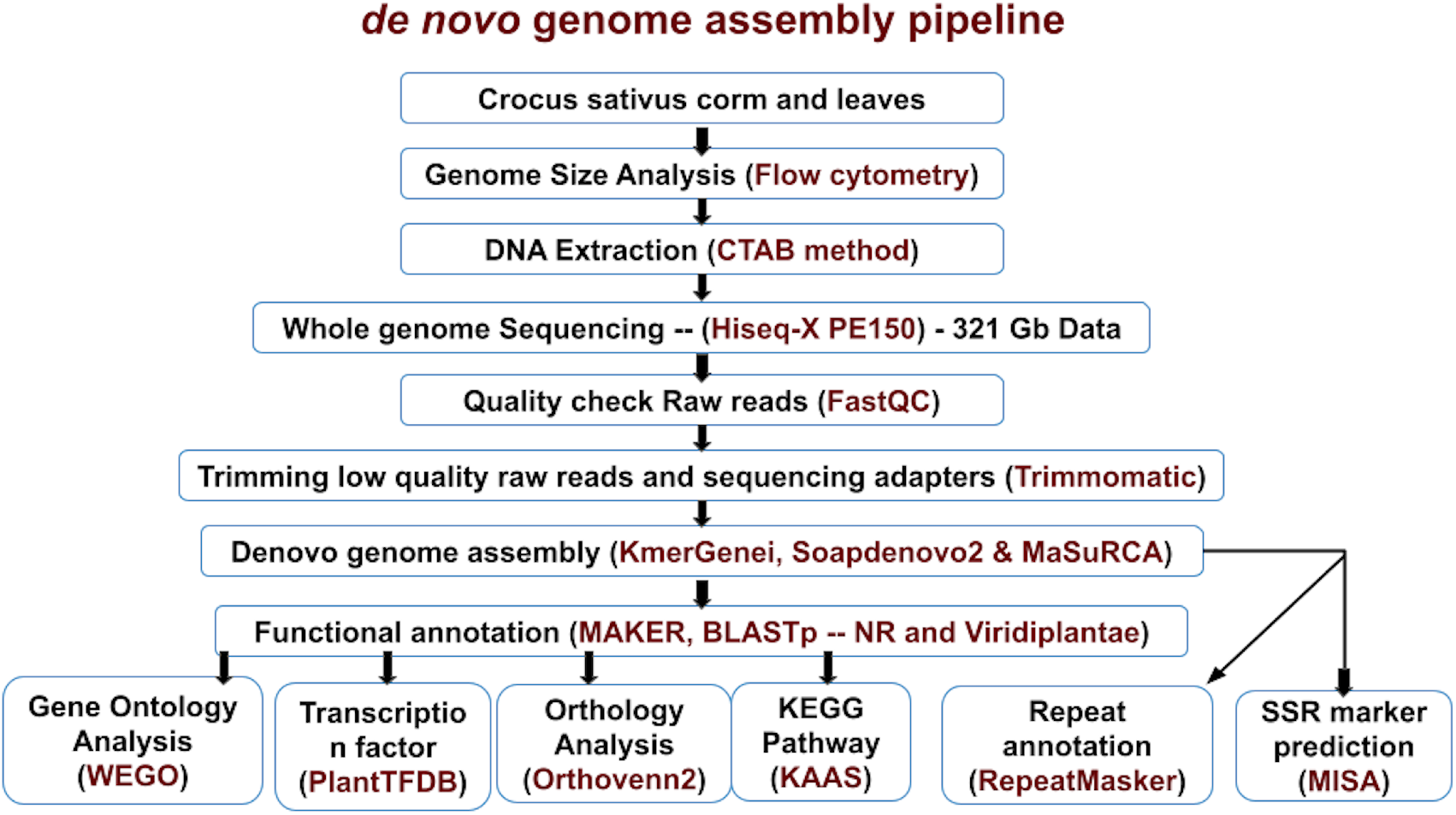
Schematic of de novo genome assembly and annotation pipeline. Black colour text represent the analytical processes and Red colour text represent the software/instrument used to perform the processes.

### *De-novo* genome assembly

*C. sativus* genome was *de-novo* assembled using SOAPdenovo2 (Luo et al., 2012) and MaSuRCA softwares (Zimin et al., 2013). Soapdenovo2 assembly with kmer 71 was comparatively better than other two kmers (69 and 73) and was designated as Cs_Assembly_1 with N50 value of 1596 and 77.9% genome coverage (Table 2). De-novo assembly with MaSuRCA was designated as Cs_Assembly_2 with N50 value of 1860 and 84.24% genome coverage. Cs_Assembly_2 was found comparatively better than Cs_Assembly_1 as the assembly statistics, such as N50, largest scaffold, genome coverage and BUSCO completeness were higher in Cs_Assembly_2 than Cs_Assembly_1. (Table 2). Further, ~87.28% of raw reads mapped back to Cs_Assembly_2, thereby indicating that most of data has been utilized for genome assembly. In addition, two previously published transcriptome data sets (Jain et al., 2016; Baba et al., 2015) were mapped to the Cs_Assembly_2 and mapping percentage of 99.92% and 92.02% were observed against Cs_Assembly_2 (Table 3). High mapping percentage represented the presence of most of the reported exons/CDS in the Cs_assembly_2 even though the genome assembly was fragmented with less N50 value. *Polygonum cuspidatum* genome was de-novo assembled using Soapdenovo2 with Illumina reads and generated an assembly of 2.56 Gb, with N50 value of 3215 and 98.5% genome coverage (Zhang et al., 2019). Similarly, the genome of *Linum usitatissimum*, flax plant was *de novo* assembled using Illumina reads having N50 scaffold of 694 Kb with 81% of genome coverage (Wang et al., 2012). Genome coverage of *C. sativus* was comparatively more than flax genome but less than *Polygonum cuspidatum* genome using same sequencing technologies.

**Table 2:** Assembly statistics of *C. sativus* genome using soapdenovo2 and MaSuRCA *de-novo* assemblers.

|  | **Soapdenovo2** |  |  | **MaSuRCA** |
| --- | --- | --- | --- | --- |
| **Assemblies** | - | **Cs_Assembly_1** | - | **Cs_Assem-**<br>**bly_2** |
| **kmers** | 69 | **71** | 73 | 99 |
| **N50 Scaffold (bases)** | 1443 | **1596** | 1508 | **1863** |
| **Number of Scaffolds** | 1537310 | **1505129** | 1433675 | **2564042** |
| **Largest Scaffold (bases)** | 45973 | **45973** | 43370 | **46734** |
| **Total sequence length** | 2684437407 | **2787926280** | 2589039086 | **3014612563** |
| **GC%** | 43.2 | **43.2** | 43.2 | **43.2** |
| **Genome Coverage (%)** | 75.01% | **77.90%** | 72.34% | **84.24%** |
| **BUSCO (%)** | 7.32% | **7.81%** | 7.05% | **44.46%** |

**Table 3:** Mapping WGS raw reads and previous published transcriptome data to Cs_assembly_2.

| **Data Type** | **WGS raw reads**<br>**in present study** | ***C.sativus***<br>**transcripts**<br>**(Jain et al., 2016)** | ***C.sativus***<br>**transcriptome raw reads**<br>**(Baba et al 2015)** |
| --- | --- | --- | --- |
| **Source of Data** | **Present study** | **Jain et al., 2016** | **Baba et al 2015** |
| **Total number of reads/transcripts** | 2135957246 | 327920 | 59043670 |
| **Mapped reads/transcripts** | 1864292472 | 327643 | 54330850 |
| **Mapping Percentage** | 87.28 % | 99.92 % | 92.02 % |

### Repeats and SSR markers

Total repeats length in *C. sativus* genome (Cs_assembly_2) was 1,460,908,750 bp (40.8%) as predicted by GenomeScope version 2. A total of 8,62,275 repeats were identified in Cs_assembly_2 wherein simple repeat (48.41%) and LTR (30.34%) were the most abundant in the genome. Specifically, Copia & Gypsy were the most abundant LTR repeats (Table 4). A total of 9,64,231 SSR markers were identified in Cs_assembly_2 wherein monomeric SSR repeats (4,86,140-50.4%) were more abundant as compared to dinucleotide (2,94,819 - 30.5%) and trinucleotide repeats (1,46,991-15.2%) with “A”, “TA”,“TTG” most abundant SSRs in each groups. The abundance of Tetranucleotide (15,375-1.59%), pentanucleotide (8596-0.9%) and hexanucleotides (12,310-1.27%) repeats each was less than 2% of total SSRs with “AAAT”, “TATAT” and “TAACCC” most abundant in respective SSRs (Table 5). SSR markers are reported to be multi-allelic, relatively abundant, widely dispersed across the genome and have been used in genetic diversity analysis, parentage assessment, species identification and mapping genetic linkage (Feng et al., 2016). These markers can be further evaluated for their application in *C. sativus*. Earlier studies on *C. sativus* transcriptome has reported the presence of 16,721 SSRs (Jain et al., 2016) and 79,028 SSRs (Qian et al., 2019) using transcriptome analysis, but higher number of SSR (9,64,231) were discovered in the present study based on genome sequence.

**Table 4:** Classification of repetitive sequences in *C. sativus* genome representing abundance of Simple repeats and LTRs.

| **Repetitive region** | **Numbers** |
| --- | --- |
| Simple repeats | 415561 |
| LTR | 260472 |
| Low_complexity | 64624 |
| DNA | 64205 |
| LINE | 45739 |
| Satellite | 3946 |
| RC_Helitron | 3072 |
| rRNA | 2749 |
| SINE | 848 |
| tRNA | 650 |
| Other | 340 |
| snRNA | 54 |
| Retroposon | 15 |
| **Total** | **862275** |

**Table 5:** SSR markers from *Crocus sativus* draft genome (Cs_assembly_2) depicting the more relative abundance of monomeric repeat microsatellite.

| SSR types | count | relative %age | Most abundant | %age |
| --- | --- | --- | --- | --- |
| monomeric repeat microsatellite | 486140 | 50.4% | “A” | 44.6 % |
| dinucleotide repeat microsatellite | 294819 | 30.5% | “TA” | 16.5 % |
| trinucleotide repeat microsatellite | 146991 | 15.2% | “TTG” | 5.72 % |
| Tetranucleotide repeat microsatellite | 15375 | 1.59% | “AAAT” | 7.7 % |
| Pentanucleotide repeat microsatellite | 8596 | 0.9% | “TATAT” | 3.2 % |
| Hexanucleotide repeat microsatellite | 12310 | 1.27% | “TAACCC” | 5.3 % |
| **Total** | **964231** |  |  |  |

### Gene prediction and annotation

In total 2,54,038 proteins were predicted from Cs_assembly_2 using MAKER pipeline. A total of 52,435 and 52,545 proteins were annotated based on BLASTp against NR and viridiplanteae database respectively (Table 6). BUSCO analysis revealed the presence of 75.7% of the plant conserved genes/orthologues in the *C. Sativus* genome. Out of total proteins, 51% (26796) were annotated to 8 top-hit plant species (Fig 2). Maximum number of proteins were annotated against *Asparagus officinalis* (9213) indicating *C. sativus* to be phylogenetically closer to *Asparagus officinalis*, as both the plants belong to same plant order Asparagales (Fig 2). 85% of total proteins (43,649) were associated with gene ontology (GO) ids and classified into biological processes (BP: 22,092 proteins) abundant in cellular and metabolic processes, cellular components (CC: 24,399 proteins) mostly localised in cell and organelle parts and molecular functions (MF: 34,442 proteins) most abundant in catalytic and transporter activities (Table S1).

**Table 6:**
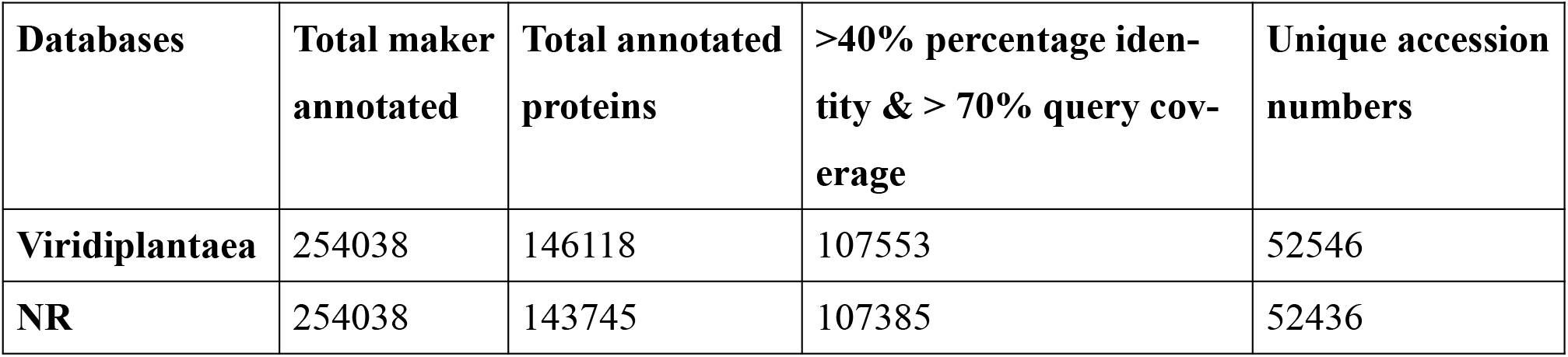
Number of genes annotated against NR and Viridiplantaea database depicting more number of proteins annotated against Viridiplantaea database.

| Databases | Total maker annotated | Total annotated proteins | >40% percentage identity & > 70% query coverage | Unique accession numbers |
| --- | --- | --- | --- | --- |
| Viridiplantaea | 254038 | 146118 | 107553 | 52546 |
| NR | 254038 | 143745 | 107385 | 52436 |

**Fig 2:**
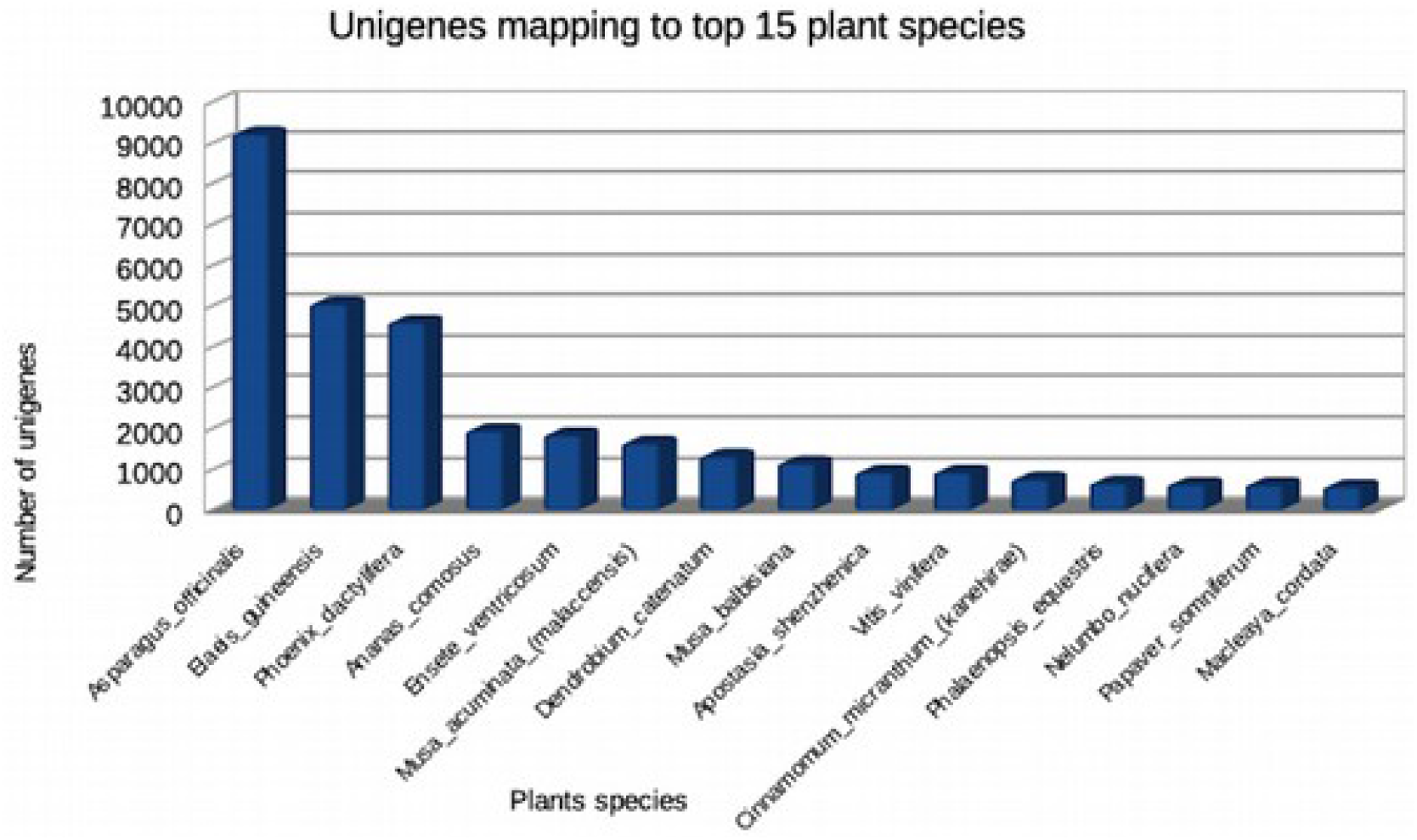
*Crocus sativus* unigenes mapping to top 15 plant species wherein most of the proteins annotated against Asparagus officinalis.

### Transcription factors

Out of the total annotated proteins, 5726 unique *C. sativus* proteins were identified as transcription factors (TFs) belonging to 57 TFs families. MYB & MYB related family proteins (11.86%), being more abundant Tfs, followed by bHLH, C2H2, NAC, FAR1, C3H, ERF, bZIP, WRKY and B3 were the top 10 abundant transcription factors family proteins (Fig 3, Table S2). TFs like MYB & MYB related, bHLH, WRKY are reported to regulate secondary metabolite (apocarotenoid) biosynthesis in *C. sativus* (Jain et al., 2016). Earlier reports on *C. sativus* transcriptome has identified less number of TFs (3819, 2601), whereas the most abundant Tfs family remains same (Jain et al., 2016; Baba et al., 2015).

**Fig 3:**
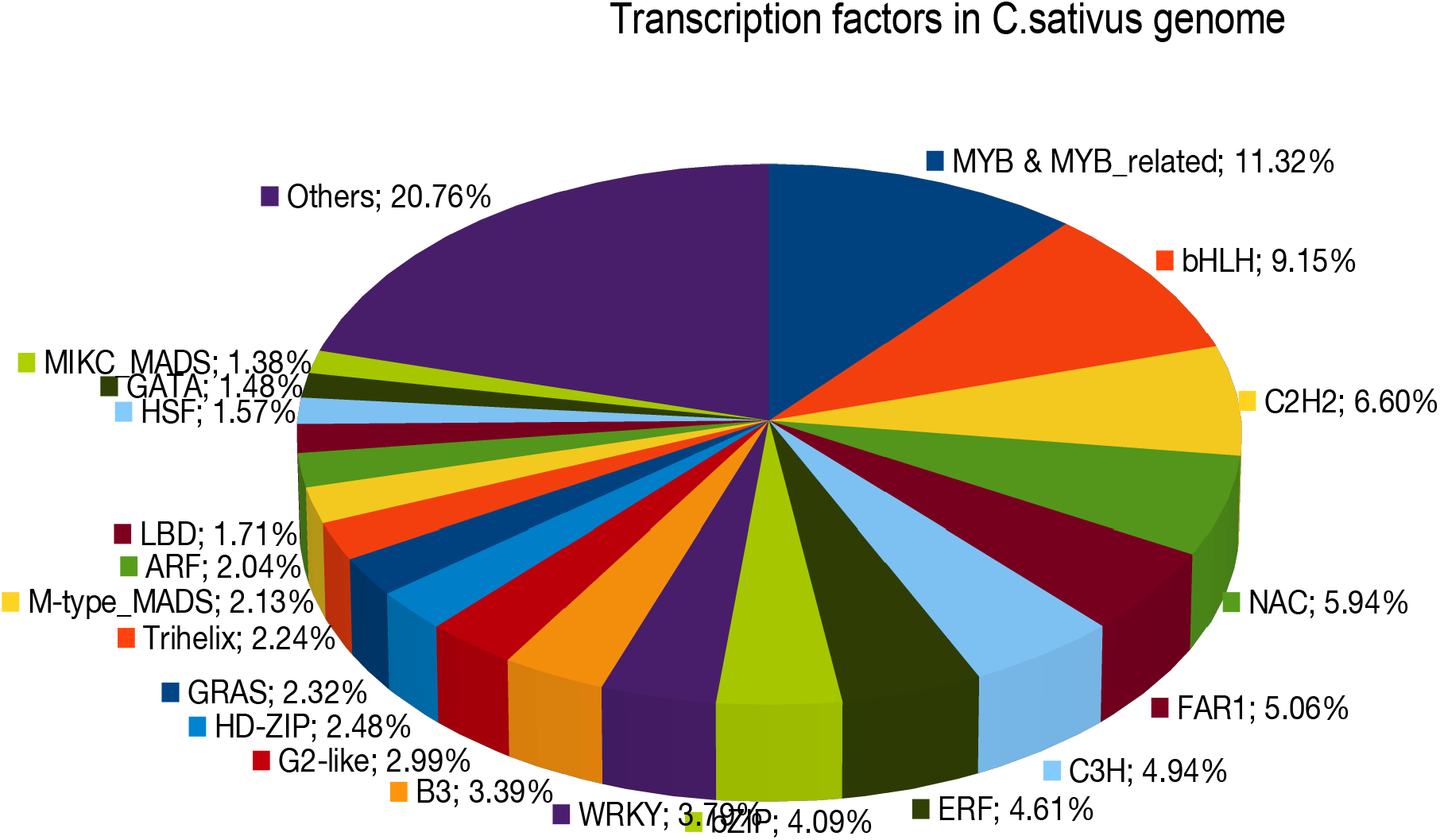
Trancription factors identified in *Crocus sativus* genome wherein MYB & MYB related Tfs were most abundant.

### Comparative genomics

*C. sativus* annotated proteins (52,545) was compared with 3 monocots plants of same order, whose genome and annotations were available in Phytozome database (David et al., 2012), namely *Asparagus officinalis, Phalaenopsis equestris, Apostatia shenzhenica* along with a model monocot plant *Oryza sativus* (Rice) using Orthovenn2. A total of 23,744 proteins cluster were found in all the plants wherein 21,606 were orthologous clusters that were atleast common in two species and 2138 were single copy gene clusters wherein each cluster have only one gene from each plant species. Conserveration of 7328 proteins clusters, comprising of 51,803 proteins, was observed among the five species (*C. sativus:10,001* proteins, *A. officinalis:9552, P. equestris:9012, A. shenzhenica:* 8570 and *O. sativa*:14,668) (Fig 4a and 4b). The conserved proteins clusters were found to be associated with biological processes (BP-23,010 proteins), cellular component (CC-582 proteins) and molecular functions (MF-957 proteins) and were enriched in defence response, RNA modification, DNA integration, regulation of transcription, rRNA processing and protein phosphorylation (Supplementary table S3). However, 2510 protein clusters (7914 proteins) were unique to *Crocus sativus* only, out of which 1636 clusters (4595 proteins) were associated with slimmed GO terms (BP: 5201, CC: 63, MF:303 proteins) associated with nucleic acid binding, transferase, hydrolase, oxidoreductase activity and protein and DNA binding activity (Supplemenatary Table S3). As per orthology analysis also, *C. sativus* was found phylogenetically closer to *A. officinalis* as more protein clusters were orthologous between *Crocus sativus* and *Asparagus officinalis* than to other plants compared in the study (Fig 5).

**Fig 4a:**
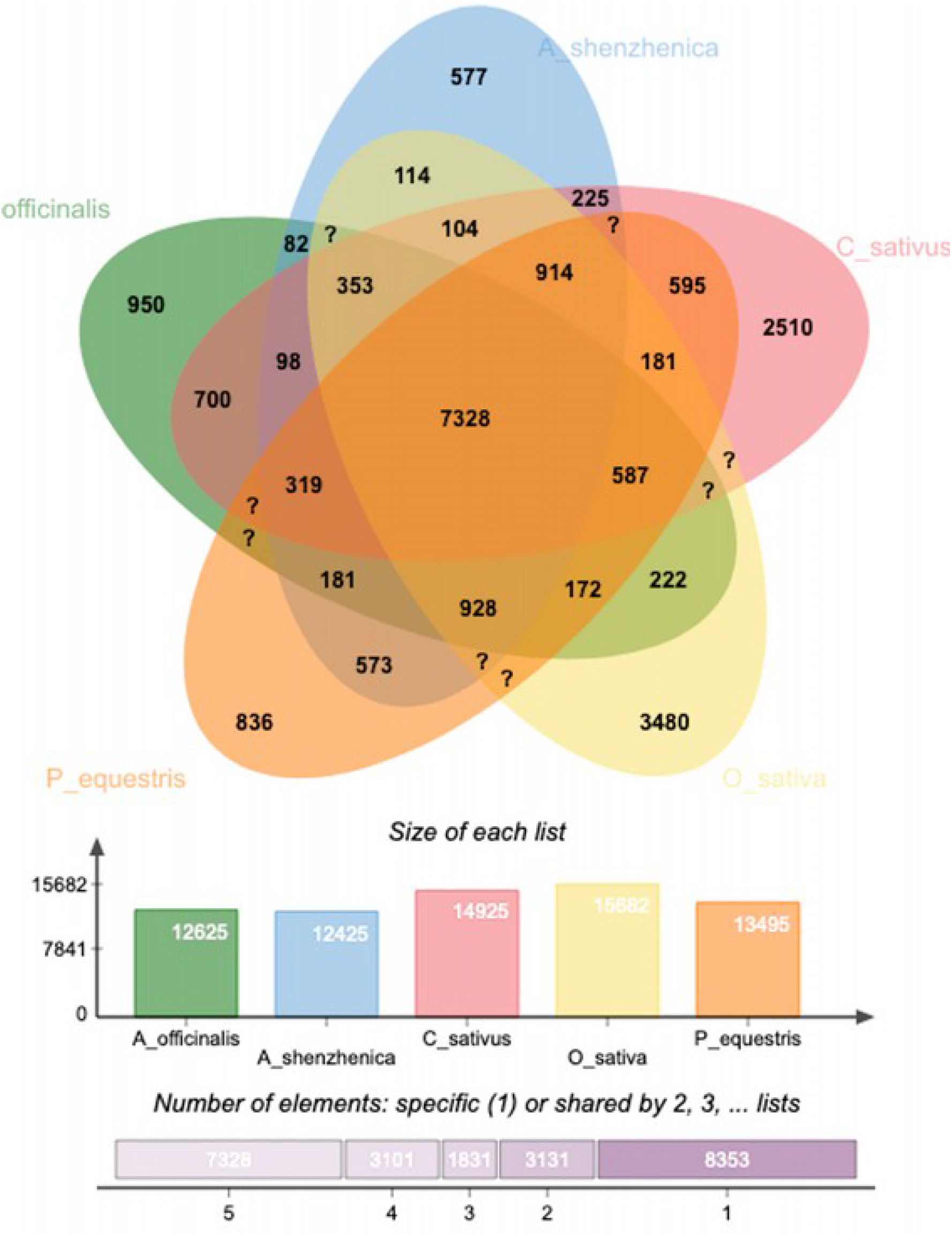
Orthology analysis of *Crocus sativus* with neighbouring plants from same order along with Rice representing 7328 proteins clusters to be conserved in all the five plant species, whereas 2510 proteins cluster were unique to *C. sativus* only.

**Fig 4b:**
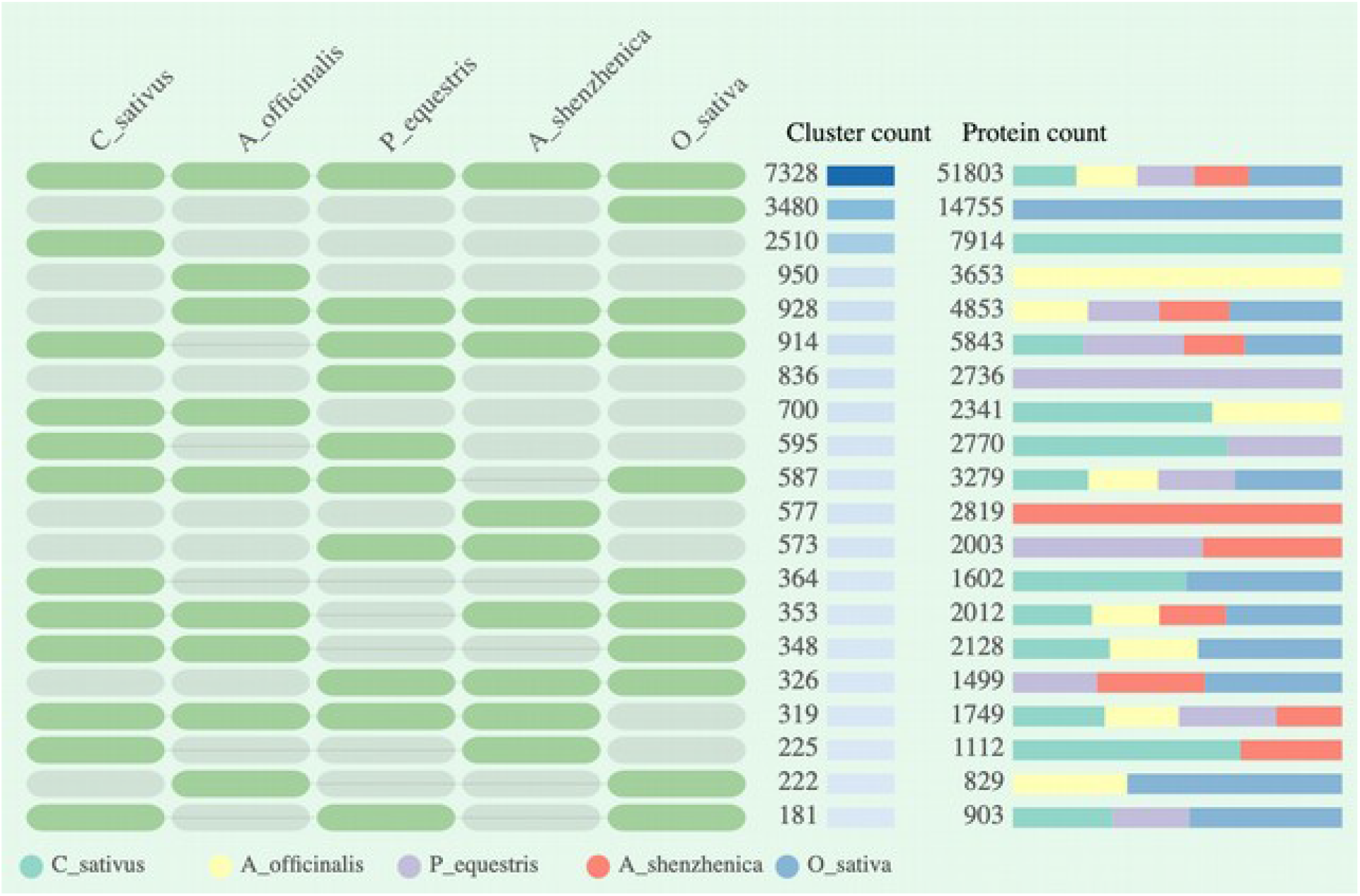
Overlapping cluster numbers between each pair of plant species representing common clusters (7328) among five plant species and unique cluster (2510) to *Crocus sativus*.

**Fig 5:**
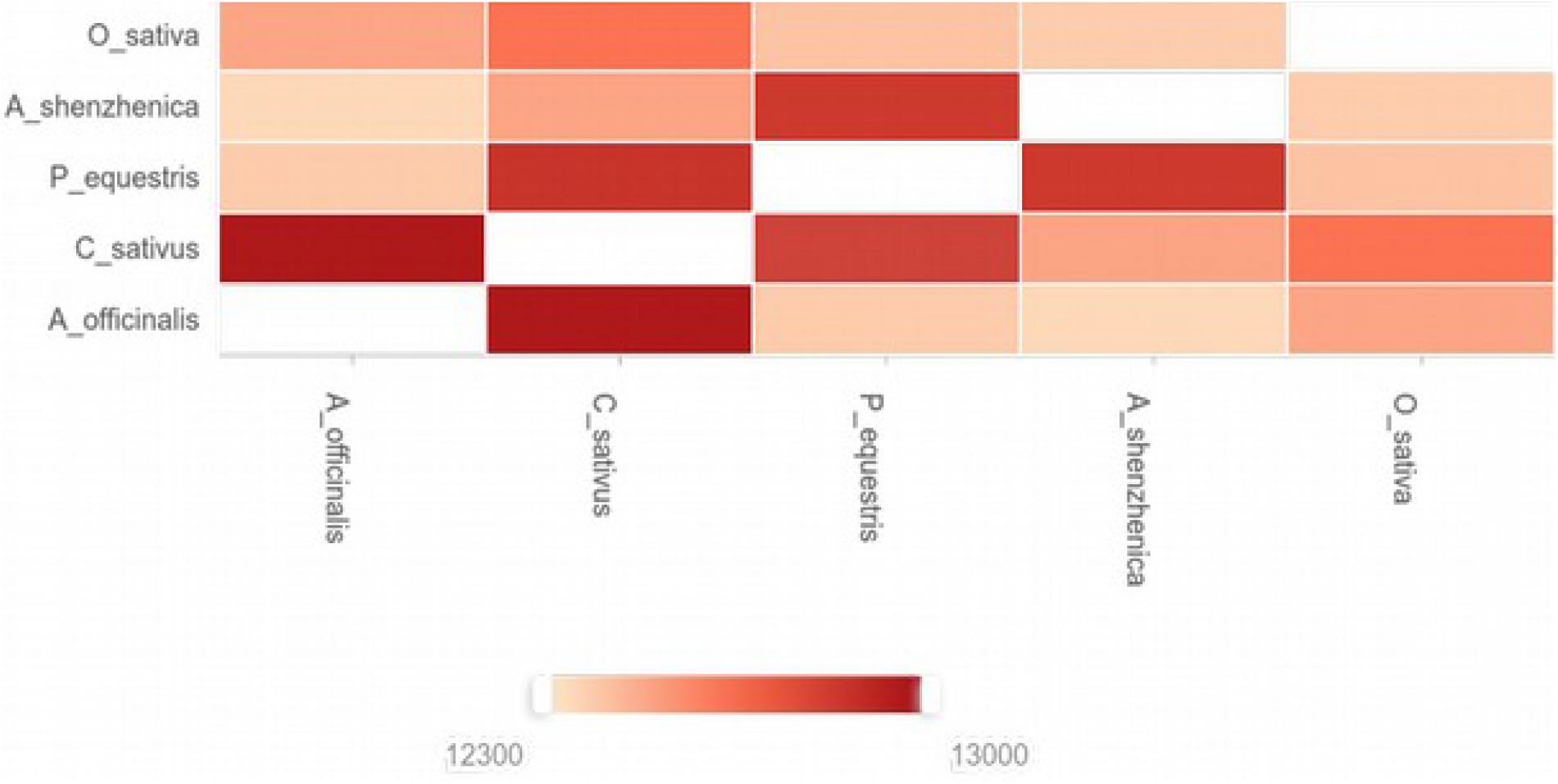
Heatmap of overlapping cluster numbers between each pair of plant species representing more number of overlapping clusters between *Crocus sativus* and *Asparagus officinalis*.

### Metabolic pathway analysis

A total of 10,912 *C. sativus* proteins were mapped to 395 KEGG pathways of monocots. Various pathways like carbohydrate metabolism, energy metabolism, lipid metabolism, nucleotide metabolism, amino acid metabolism, glycan metabolism, metabolism of cofactors and vitamins along with biosynthesis of terpenoids, polyketides and other secondary metabolites were found complete wherein all the genes involved in pathway were present in draft assembly (Table S4). We, further, investigated the presence of genes involved in the synthesis of apocarotenoids namely crocins, picrocrocin, and safranal that are produced in the stigma of *C. Sativus*. These apocarotenoids impart red color, bitter taste, and pungent aroma to stigma of saffron and have various medicinal properties (Maggi et al., 2020). The molecular basis of apocarotenoid biosynthesis in *C. Sativus* has been well studied using transcriptomics studies (Baba et al., 2015; Jain et al., 2016). In the present study, the genes encoding the enzymes involved in carotene biosynthesis pathway, regulating the apocarotenoids synthesis, were present in the *C.sativus* genome (Fig 6). This is the first de novo draft genome sequence of *Crocus sativus* that needs to be complemented with the long read sequencing technology (PacBio) to fill in the gaps in the present genome to generate a complete genome sequence. However this draft genome sequence, in addition to revealing previous unknown genomic information on saffron, will also be used as a reference genome for future genome sequencing attempts in saffron.

**Fig 6:**
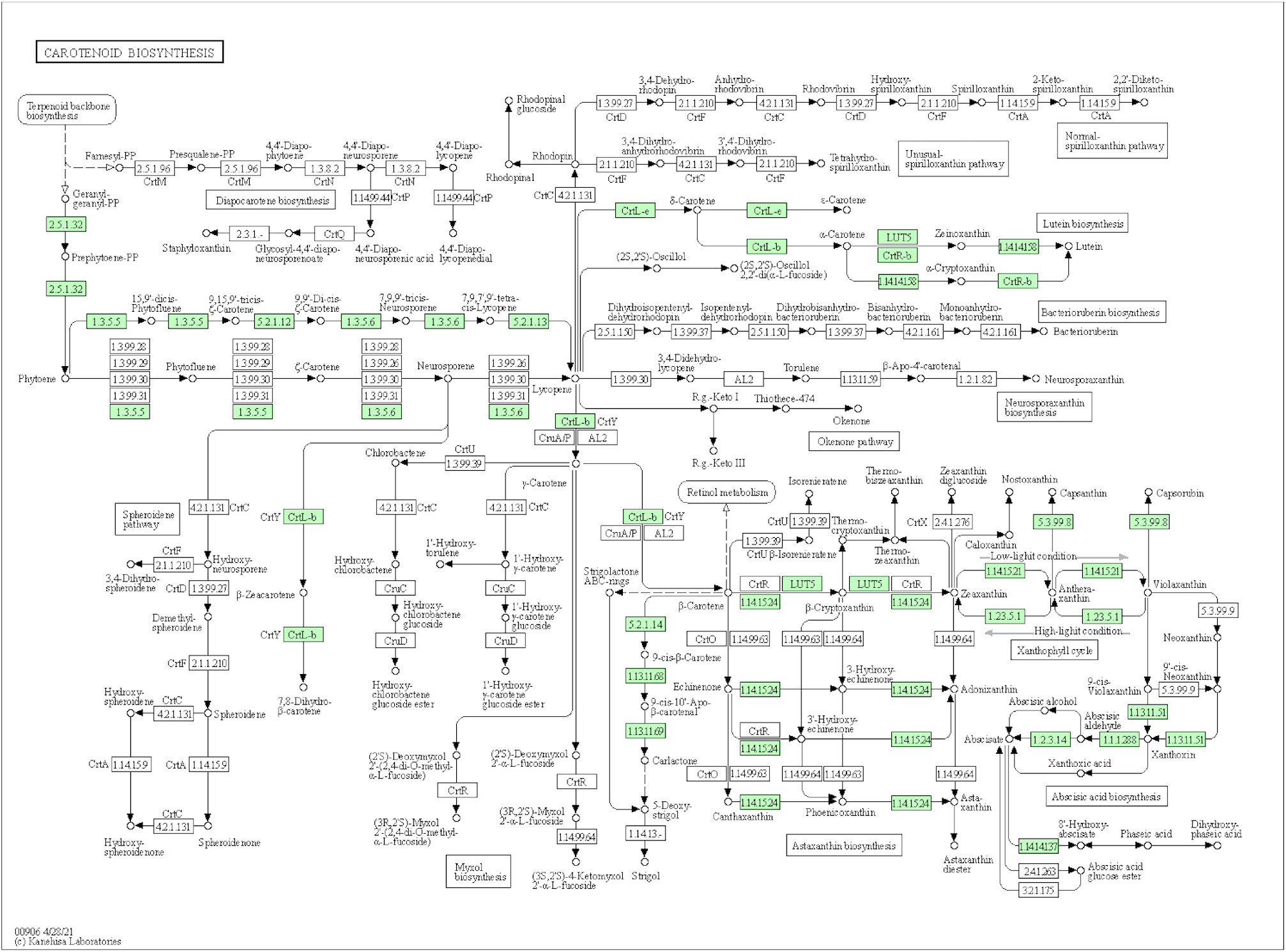
Carotene biosynthesis pathway in *Crocus sativus* that was found complete with all the genes in the pathway.

## Acknowledgments

Authors are grateful to Mr. Shanu Magotra, School of Biotechnology, University of Jammu, J&K for his help in sample collection from Kistwar, Jammu and Kashmir. Authors are thankful to NCBS CIFF facility for their help in flowcytometry analysis. SA is thankful to DST-Women Scientist-A research grant (SR/WOS-A/LS-96-2018) for funding this research.

